# Radiochemical Synthesis and Evaluation of 3-[^11^C]Methyl-4-aminopyridine in Rodents and Non-Human Primates for Imaging Potassium Channels in the CNS

**DOI:** 10.1101/2022.06.22.495540

**Authors:** Yang Sun, Nicolas J. Guehl, Yu-Peng Zhou, Kazue Takahashi, Vasily Belov, Maeva Dhaynaut, Sung-Hyun Moon, Georges El Fakhri, Marc D. Normandin, Pedro Brugarolas

## Abstract

Demyelination, the loss of the insulating sheath of neurons, causes failed or slowed neuronal conduction and contributes to the neurological symptoms in multiple sclerosis, traumatic brain and spinal cord injuries, stroke, and dementia. In demyelinated neurons, the axonal potassium channels K_v_1.1 and K_v_1.2, generally under the myelin sheath, become exposed and upregulated. Therefore, imaging these channels using positron emission tomography can provide valuable information for disease diagnosis and monitoring. Here, we describe the novel tracer for K_v_1 channels [^11^C]3-methyl-4-aminopyridine ([^11^C]3Me4AP). [^11^C]3Me4AP was efficiently synthesized via Pd(0)-Cu(I) co-mediated Stille cross-coupling of a stannyl precursor containing a free amino group. Evaluation of its imaging properties in rats and nonhuman primates showed that [^11^C]3Me4AP has a moderate brain permeability and slow kinetics. Additional evaluation in monkeys showed that the tracer is metabolically stable and that a 1-tissue compartment model can accurately model the regional brain time-activity curves. Compared to the related tracers [^18^F]3-fluoro-4-aminopyridine ([^18^F]3F4AP) and [^11^C]3-methoxy-4-aminopyridine ([^11^C]3MeO4AP), [^11^C]3Me4AP shows lower initial brain uptake, which indicates reduced permeability to the blood-brain-barrier and slower kinetics, suggesting higher binding affinity consistent with *in vitro* studies. While the slow kinetics and strong binding affinity resulted in a tracer with less favorable properties for imaging the brain than its predecessors, these properties may make 3Me4AP useful as a therapeutic.

## Introduction

Myelin is the protective sheath that surrounds most axons in the brain and spinal cord and speeds up impulse propagation along the axons of neurons. Demyelination occurs when the myelin is damaged from various diseases, including multiple sclerosis (MS)^1^, brain and spinal cord injury^2, 3^, stroke^4^ and Alzheimer’s disease^5^. However, the time course of demyelination and its contribution to disease symptoms and progression remains unclear. Positron emission tomography (PET) imaging has the potential to detect and quantify biochemical processes underlying demyelination and thus serve for monitoring the progress of remyelinating therapies^6, 7^. Following demyelination, the axonal voltage-gated potassium channels K_v_1.1 and K_v_1.2, normally buried beneath the myelin sheath, become exposed and increase in expression^8-12^. Based on the clinically approved MS drug 4-aminopyridine (4AP), our group previously proposed a radiofluorinated analog of 4AP (3-[^18^F]fluoro-4-aminopyridine, [^18^F]3F4AP) that could be used to image demyelination. This hypothesis was confirmed experimentally in rodent models of MS^13^. Further evaluation of [^18^F]3F4AP in non-human primates (NHPs) showed that [^18^F]3F4AP has excellent properties for primate brain imaging, such as high brain penetration, fast kinetics, and high metabolic stability. Furthermore, [^18^F]3F4AP was very sensitive to a focal brain injury in a monkey^14^. More recently, due to the advantages of having short-lived radiotracers, such as allowing multiple scans on the same subject on the same day, we explored 3-[^11^C]trifluoromethyl-4-aminopyridine ([^11^C]3CF_3_4AP)^15^ and 3-[^11^C]methoxy-4-aminopyridine ([^11^C]3MeO4AP)^16^ as viable ^11^C-labeled alternatives. Compared to [^18^F]3F4AP, [^11^C]3CF_3_4AP showed higher brain penetration and faster washout with lower heterogeneity across brain regions, which is consistent with its higher lipophilicity and lower binding affinity as was measured *in vitro*^17^. Comparatively, [^11^C]3MeO4AP showed lower initial brain penetration and slower brain kinetics than [^18^F]3F4AP, indicative of its lower lipophilicity and higher binding affinity. A quantitative comparison of [^11^C]MeO4AP and [^18^F]3F4AP showed a high correlation in volumes of distribution (V_T_) across brain regions, suggesting that these compounds bind to the same target. A recent detailed study of four novel 4-aminopyridine potassium channel blockers by electrophysiology showed that 3-methyl-4-aminopyridine (3Me4AP) has a higher binding affinity than 4AP, 3F4AP, 3CF_3_4AP and 3MeO4AP^17^. Moreover, measurement of the basicity (p*K*_a_) and lipophilicity (logD) of these K^+^ channel blockers showed that 3Me4AP is more basic (higher p*K*_a_) than 4AP, 3MeO4AP, 3CF_3_4AP and 3F4AP and more polar (lower logD) than all these compounds except 4AP. These counterbalanced properties (high binding affinity and high polarity) make 3Me4AP an attractive candidate tracer for imaging potassium channels. In the present work, we describe the radiochemical synthesis of [^11^C]3Me4AP and the evaluation of its imaging properties in rodents and non-human primates. We also perform a detailed comparison between [^18^F]3F4AP, [^11^C]MeO4AP and [^11^C]Me4AP.

## Results and Discussion

### Radiochemical synthesis of 3-[^11^C]methyl-4-aminopyridine ([^11^C]3Me4AP) by Pd(0)-Cu(I) co-mediated ^11^C-methylation

The most common ^11^C-labeling strategy for the synthesis of ^11^C-labeled PET tracers is ^11^C-methylation *via* a nucleophilic substitution with [^11^C]methyl iodide (MeI) or triflate (MeOTf)^18-20^. However, this method is only suitable for methylating heteroatom nucleophiles to form O–[^11^C]CH_3_, N–[^11^C]CH_3_, and S–[^11^C]CH_3_^(17,18)^. Several palladium-mediated Suzuki and Stille cross-coupling reactions of [11C]MeI with the corresponding organoboron and organostannyl precursors have been reported in the last decade, allowing a direct C–[^11^C] bond formation^21^. In addition, copper(I) salt has been used in combination with organostannyl precursors to enhance the cross-coupling by generating *in situ* an organocuprate reagent^22^. Such intermediate reduces the energy barrier of the transmetalation, which is the rate-determining step in the catalytic cycle of the Stille reaction. Despite these advances, multiple-step synthesis^23^ or harsh conditions^(18)^, such as high temperature and microwave, are required to achieve satisfying efficiency. Based on these previous reports, we hypothesized that 3-(tributylstannyl)pyridin-4-amine **2** would be a good precursor for our desired target compound [^11^C]3Me4AP **3**, considering the stability and functional group tolerance of tributylstannyl precursors in cross-coupling reactions^24^. Consequently, **2** was prepared from 3-iodopyridin-4-amine **1** without pre-protection in one pot via sequential deprotonation, lithium-halogen exchange and stannylation in a 49% isolated yield (**Figure 1A, Equation 1**). Next, single-step labeling of **2** with [^11^C]MeI *via* the Pd(0)-Cu(I) co-mediated reaction gave the desired [^11^C]3Me4AP in an overall 89 ± 5% (n = 5) decay corrected radiochemical yield by analytical HPLC with the only byproduct being unreacted [^11^C]MeI (see **Figure 1B**). The product was purified using normal phase semipreparative HPLC. On semiprep HPLC, the product peak was broad, and we typically only collected the tip of the peak resulting in an average of 51 ± 4% (n=4) decay-corrected isolated yield (prep-HPLC see **Figure 1C**). The molar activity was excellent (up to 129.5 GBq/μmol), and so was the radiochemical purity (>99%) (**Figure 1A**, Equation 2). The identity of [^11^C]3Me4AP was confirmed by analytical HPLC (**Figure 1D-E**). Avoiding the typical protection and deprotection steps of the amino group reduced the synthesis time, which is critical when working with the relatively fast decaying radionuclide carbon-11 (*t*_1/2_ = 20.3 min). From the end of the bombardment to reformulation, it took about 50 min.

**Figure 1.**
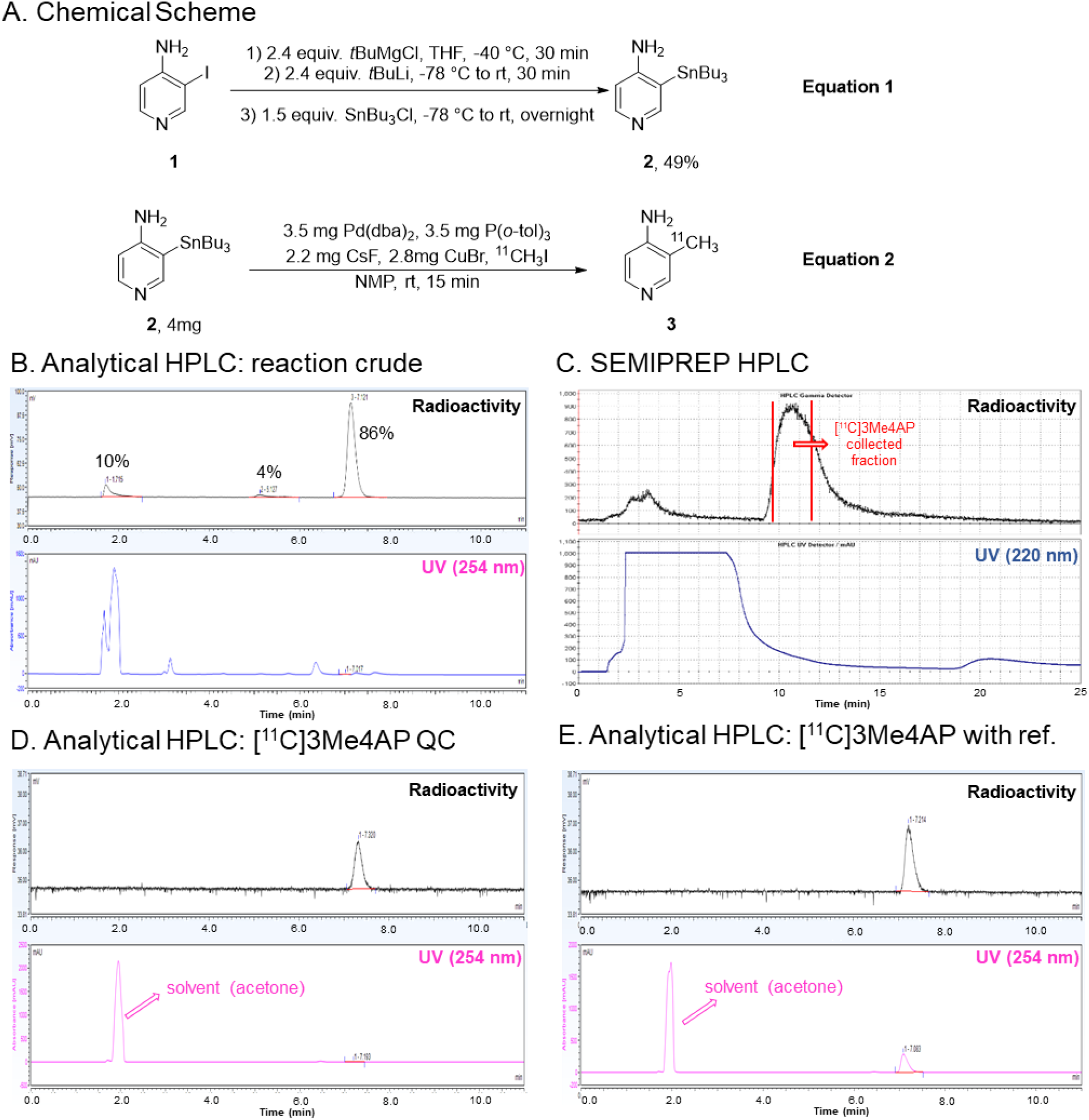
Radiochemical synthesis of [^11^C]3Me4AP. (A) chemical reaction scheme of precursor preparation and radiosynthesis of [^11^C]3Me4AP; (B) analytical HPLC (radio and 254 nm UV chromatograms) of the crude reaction; (C) semipreparative HPLC (high-performance liquid chromatography) purification (radio and 220 nm UV chromatograms); (D) analytical HPLC (radio and 254 nm UV chromatograms) of tracer only; (E) analytical HPLC (radio and 254 nm UV chromatograms) of tracer plus reference standard.

### [^11^C]3Me4AP in healthy rats: PET imaging, time-activity curves and biodistribution

To evaluate the potential use of [^11^C]3Me4AP as a PET tracer for brain imaging, PET imaging in healthy female and male rats was conducted following intravenous tail injection of [^11^C]3Me4AP (1.5 to 2.7 μCi/g) alone or [^11^C]3Me4AP (same) plus 5 mg/kg of non-radiolabeled 3Me4AP (coinjection). The summed PET images from 10 to 60 min showed high uptake in the thyroid and salivary glands (**Fig. 2A**), similar to what has been observed with [^18^F]3F4AP. Radioactivity in the brain peaked early (within the first 3 min) and was followed by a short phase where the activity increased (3-10 min) and a very slow washout (**Fig. 2B**). The brain time-activity curves (TACs) after coinjection of the tracer with 5 mg/kg of 3Me4AP showed a similar initial brain uptake and a higher signal after 10 min. To confirm the higher brain uptake, additional animals were injected with [^11^C]3Me4AP with or without the cold compound, and the radioactivity in the brain and blood was measured *ex vivo* using a gamma counter. At 60 min, the whole brain to blood standardized uptake value ratio (SUVR) was 43% higher in the coinjection group than in the baseline group (2.65 ± 0.67, n = 8, *vs*. 1.85 ± 0.23, n = 7, *p* = 0.017) (**Fig. 2C**). This increase in brain uptake upon coinjection of cold tracer is consistent with previous reports describing that high doses of 4AP analogs cause more channels to transition to the open (bindable) conformation and thus result in higher receptor binding^14, 25^.

**Figure 2.**
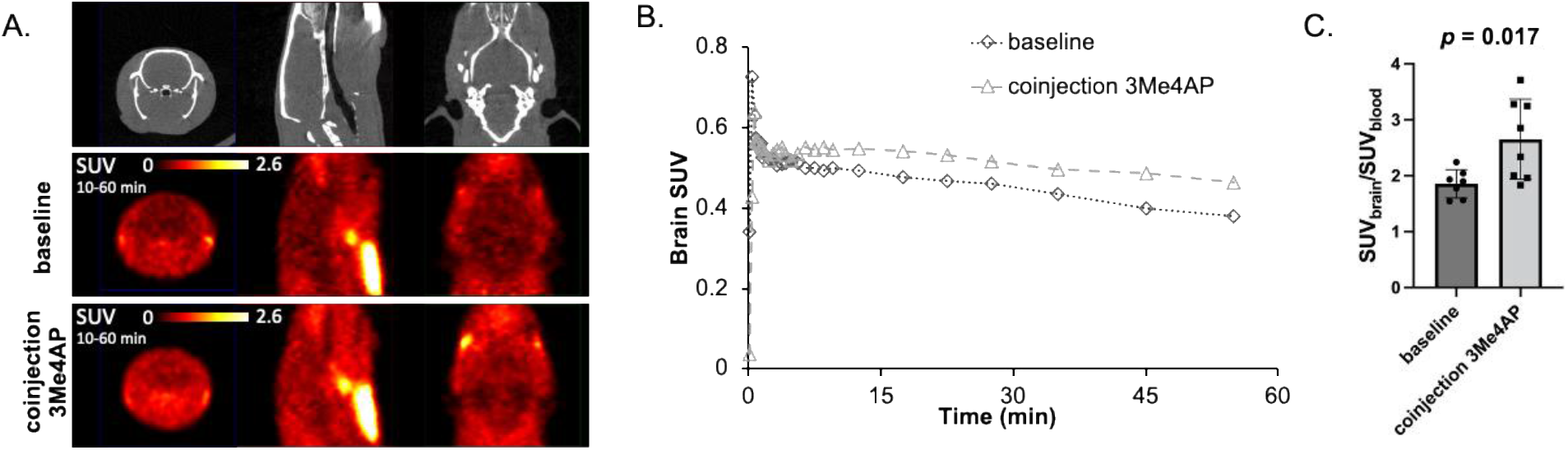
[^11^C]3Me4AP in rats. (A) Representative CT and PET images of two rats: one injected with [^11^C]3Me4AP only (baseline) and the other injected with [^11^C]3Me4AP plus 5mg/kg of cold 3Me4AP. (B) Brain time-activity curves (TACs) of the animals shown in A. (C) *Ex vivo* SUV rat blood and brain 60 min after injection of [^11^C]3Me4AP and [^11^C]3Me4AP plus 5-10 mg/kg of cold 3Me4AP. Each dot represents a different animal.

### [^11^C]3Me4AP in Non-Human Primates: Brain Uptake and Pharmacokinetic Modeling

Given the promising results obtained in rodents, dynamic PET imaging with arterial blood sampling was performed in two non-human primates after injection of [^11^C]3Me4AP. Each animal was scanned twice under baseline conditions.

#### Analysis of [^11^C]3Me4AP in blood

Radioactivity time courses in whole-blood (WB) and plasma (PL) were consistent between the two scans from M2 and the first scan in M1 (**Fig. 3A,B**). The second scan in M1, however, showed slightly faster blood clearance (**Sup. Fig. 1**). Plasma free fraction measurements performed in triplicate for each scan indicated that [^11^C]3Me4AP is mostly free in plasma (0.94 ± 0.03, N=4) with no differences between animals. WB to PL radioactivity concentration ratio reached a plateau within 1 min (WB/PL ratio at 1 min postinjection across all studies = 1.28 ± 0.01, **Fig. 3C**). RadioHPLC measurements demonstrated high *in vivo* stability with ∼90% of activity attributable to [^11^C]3Me4AP even after 90 min (**Fig. 3D,E**). Due to the moderately low signal-to-noise ratio (SNR) observed in the metabolite measurements of plasma samples drawn toward the end of the scan, metabolite measurements were averaged across the two baseline scans within each animal in order to derive the metabolite-corrected input function used in the subsequent pharmacokinetic modeling analysis.

**Figure 3.**
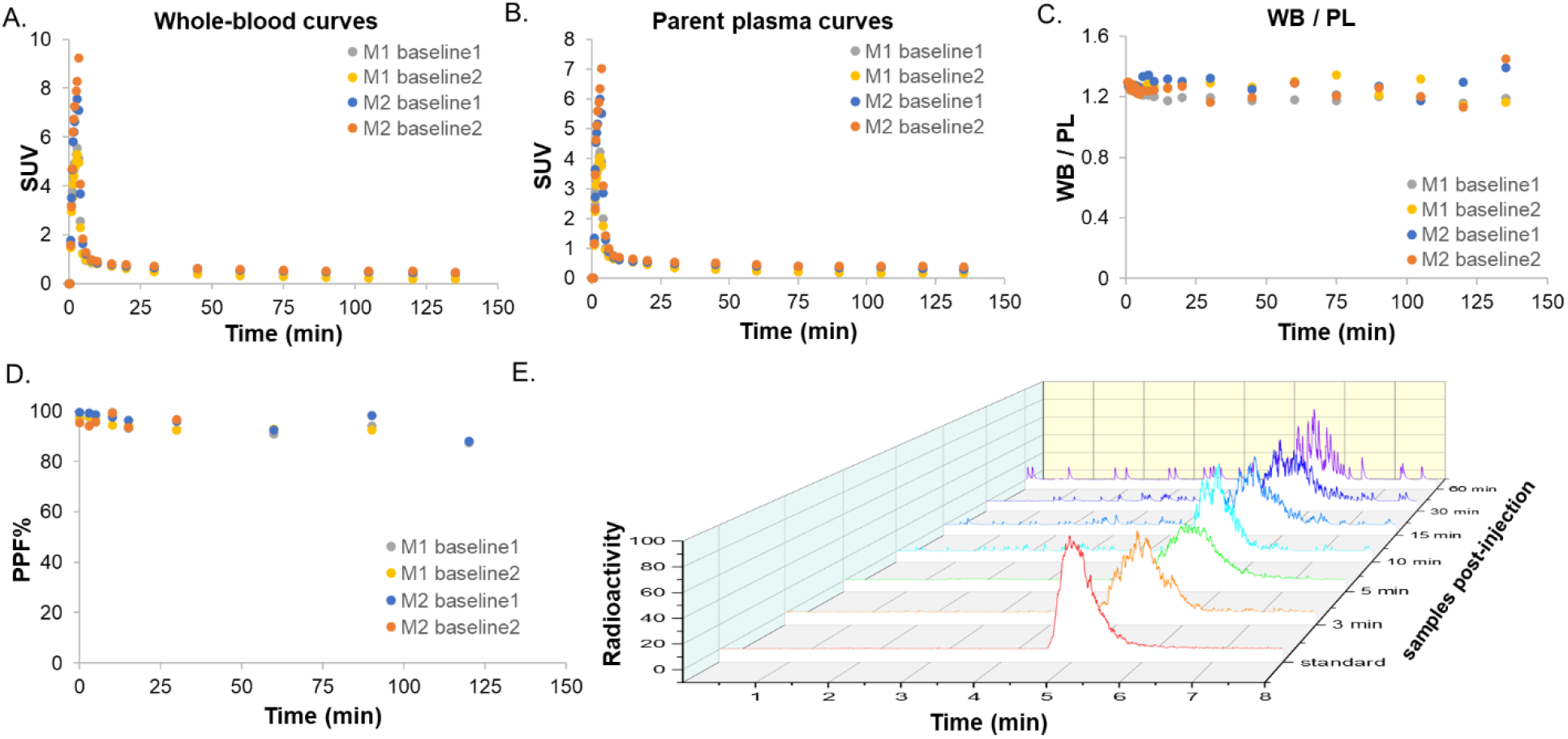
[^11^C]3Me4AP in arterial blood. (A) Time course in whole blood (WB); (B) Time course in plasma (PL); (C) Time course of WB to PL radioactivity concentration ratio; (D) [^11^C]3Me4AP percent parent in plasma (%PP) in blood samples drawn up to 120 min post tracer administration. (E) Representative radiochromatograms of arterial plasma samples drawn up to 60 min post tracer administration. In the legends, M1 and M2 refer to monkey 1 and monkey 2, respectively.

#### Analysis of [^11^C]3Me4AP in brain

In the monkey brain, [^11^C]3Me4AP showed moderate initial brain penetration (whole brain SUV at 3 min = 0.7) followed by a slow increase in brain uptake. Brain time activity curves (TACs) were largely consistent in three of the four scans (**Sup. Fig. 2**). The second scan in M1, however, showed more reversible [^11^C]3Me4AP brain kinetics, which was consistent with faster blood clearance observed in this particular study. Summed images from 60 to 120 min (**Fig. 4A**) and brain TACs (**Fig. 4B**) showed moderate heterogeneity across brain regions, with the highest uptake in the cortex and lowest in the white matter. Pharmacokinetic analysis was performed using the full scan duration (150 min) as well as after truncating the data to 120, 90 and 60 min. Visual inspection of the model fits demonstrated good fits in all brain regions and for all scans using a one-tissue (1T) compartment model with the vascular contribution to the PET signal included as a model parameter (1T2k1v). The Akaike Information Criterion (AIC)^26^ used to assess the goodness of fits across alternative models confirmed that the 1T2k1v was the preferred model. Overall, *V*_T_ estimates obtained while truncating the scan duration were in good agreement with those obtained while using the full 150 minutes of data demonstrating good stability of *V*_T_ measurements (**Table 1**). Most of the variability in the *V*_T_ estimates can be attributed to variation in *V*_T_ measurements in small brain regions like the amygdala and hippocampus, which presented higher noise level, especially towards the end of the scan (**Figure 4C**). Using 120 min of data provided the best reproducibility (**Table 1**) between baseline scans compared to using other scan durations. However, due to the slow kinetics of this tracer, 150 minutes of data was selected as the reference duration for the remaining analyses performed in this work.

**Figure 4.**
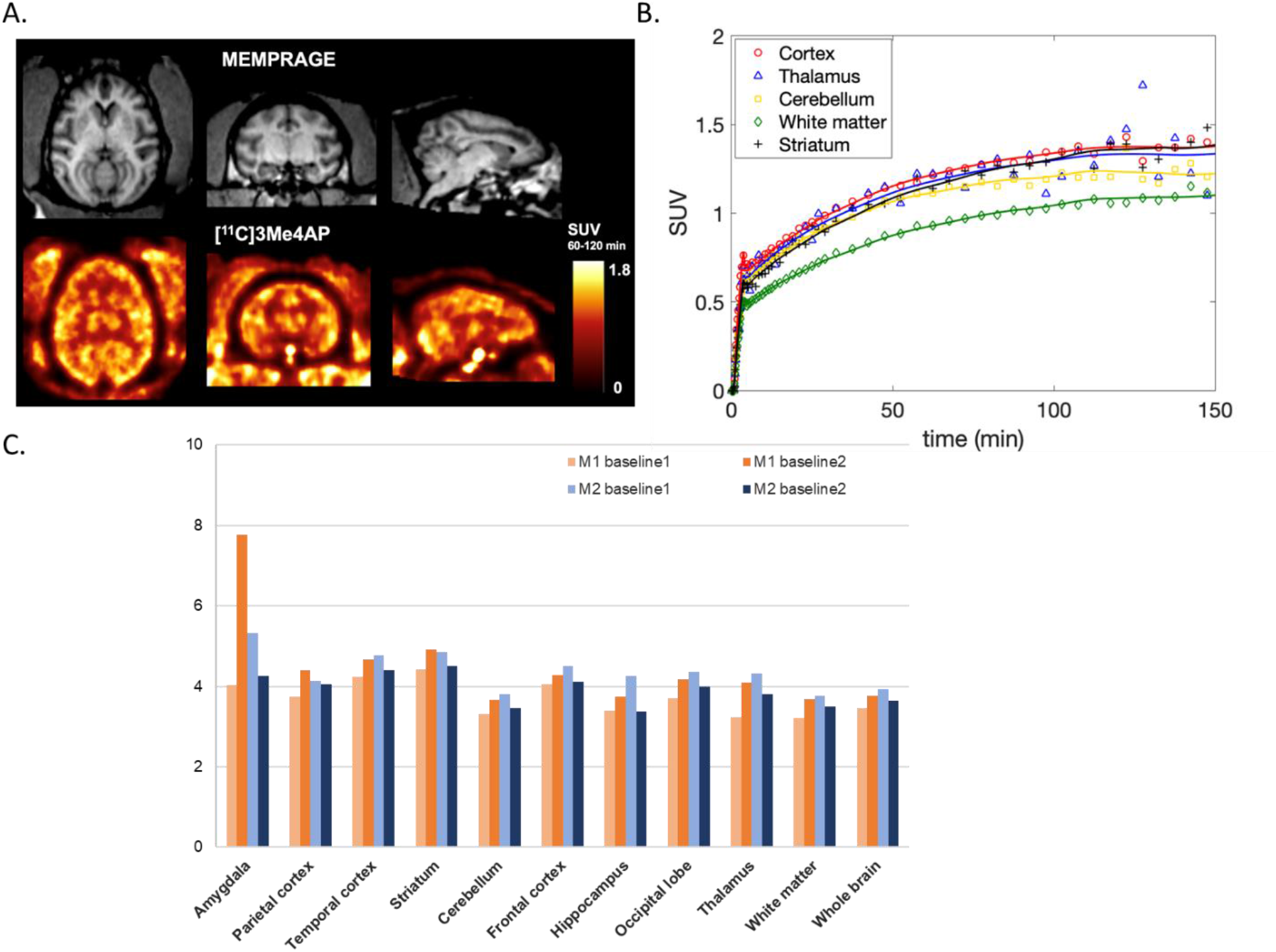
[^11^C]3Me4AP in monkey brain. (A) Representative MRI MEMPRAGE (top) and representative PET SUV images calculated from 60-120 min post-injection (bottom) of M2. Images are presented in the MRI NIMH macaque template (NMT) space. (B) Time activity curves of selected brain regions in the same animal. (C) Regional 1T2k1v VT Estimates (mL/cc) for each baseline scan and animal.

**Table 1.** Time stability (relative to full scan duration) and reproducibility of *V*_*T*_ estimates between baseline scans using different PET data truncations. Data are presented for each animal as average ± standard deviation across all brain regions.

| Data duration | Time stability (%) | Test-retest M1 (%) | Test-retest M2 (%) |
| --- | --- | --- | --- |
| 0-150 min | NA | 17.4 ± 16.9 | 11.0 ± 6.7 |
| 0-120 min | -0.6 ± 7.5 | 12.7 ± 9.8 | 7.3 ± 6.0 |
| 0-90 min | 1.2 ± 12.2 | 20.6 ± 20.6 | 10.6 ± 7.3 |
| 0-60 min | 0.5 ± 20.0 | 25.4 ± 31.0 | 17.0 ± 8.9 |

*K*_*1*_ estimates – representing the rate constant for transfer of tracer from plasma to the brain tissue – quantitatively confirmed moderate brain penetration (*K*_*1*_ in whole brain ∼0.035 mL/min/cc). The total volume of distribution (*V*_*T*_) values ranged from 3.61 mL/cc and 3.78 mL/cc in the white matter to 4.67 mL/cc and 4.68 mL/cc in the striatum of Monkey 1 and 2, respectively (**Figure 4c**).

In addition to compartmental modeling, the Logan and MA1 graphical methods were tested for the estimation of *V*_*T*_. The *V*_*T*_ estimates obtained with the Logan graphical method underestimated compartmental *V*_*T*_ values (mean difference = -11.8 ± 13.1% across brain regions using a *t\** of 20 min), and this underestimation was exacerbated in the small brain regions surveyed in this work such as in amygdala (-45.0 ± 10.9%) and hippocampus (-19.7 ± 13.6%). This underestimation was likely due to the increased noise level in those small regions combined with the slow kinetics of [^11^C]3Me4AP in the monkey brain, as previously reported for the Logan graphical method with other tracers. ^27^ On the other hand, *V*_*T*_ estimates obtained with MA1 and using a *t\** of 20 min demonstrated a good agreement with *V*_*T*_s obtained from the 1T model (*V*_*T*,MA1_=0.92 *V*_*T*,1T2k1v_+0.18; r=0.99, p < 0.0001; mean difference= -3.2±2.3% across brain regions and studies).

#### Quantitative comparison of [^11^C]3Me4AP with [^18^F]3F4AP and [^11^C]3MeO4AP

Prior comparison between the previously developed tracers [^11^C]3MeO4AP and [^18^F]3F4AP using the Guo method^28^ had shown a very good correlation between the two tracers (*r* = 0.93), suggesting a common target^16^. This comparison also indicated that [^11^C]3MeO4AP has a higher binding affinity and specific binding than [^18^F]3F4AP. In the present work, a comparison between the *V*_*T*_ estimates from [^11^C]3Me4AP and [^18^F]3F4AP showed a weaker linear correlation (**Fig. 5A**, *r* = 0.75) which, although it still suggests a common target, may be indicative of higher variability in the *V*_*T*_ estimates. The correlation plot showed a slope greater than 1 (1.93, p = 0.0001 for both monkeys combined), which suggests that [^11^C]3Me4AP has a higher *in vivo* binding affinity than [^18^F]3F4AP which is consistent with the dissociation constants measured using electrophysiology^17^. When comparing [^11^C]3Me4AP with [^11^C]3MeO4AP, the linear correlation in *V*_*T*_ values was also moderate (**Fig. 5B**, *r* = 0.79). In this comparison, the slope of the linear regression was smaller than 1 (0.73, p < 0.0001 for both monkeys combined), suggesting that [^11^C]3Me4AP has slightly lower *in vivo* affinity than [^11^C]3MeO4AP. The Guo plot can also provide information about the relative specific binding of the two compounds being compared by looking at the sign of the intercept. When [^11^C]Me4AP was compared with [^18^F]3F4AP, the intercept value was close to 0 (y = 0.10, p = 0.91). and when compared with [^11^C]3MeO4AP the intercept was greater than 0 (y = 0.67, p = 0.31). However, given the high p-values, it is not possible to draw meaningful conclusions regarding the relative specific binding.

**Figure 5.**
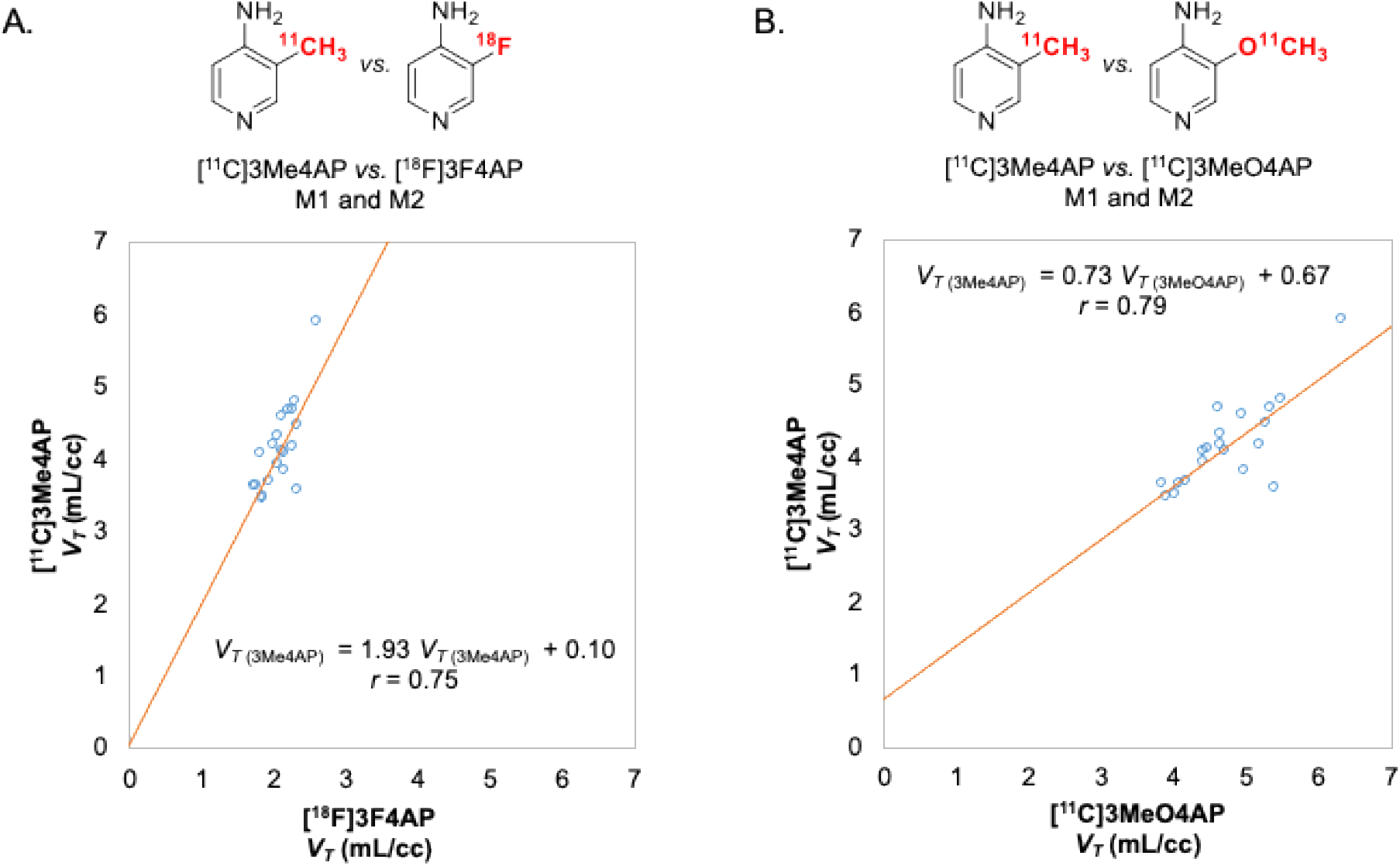
Comparison of [^11^C]3Me4AP with [^18^F]3F4AP and [^11^C]3MeO4AP. **A**. [^11^C]3Me4AP *vs*. [^18^F]3F4AP. **B**. [^11^C]3Me4AP *vs*. [^11^C]3MeO4AP.

One more aspect worth noting is that one of the monkeys used in these studies (M2) had a minor focal lesion in the brain. This lesion could be easily visualized and quantified with [^18^F]3F4AP and [^11^C]3MeO4AP and displayed a higher *V*_*T*_ value and lower *K*_*1*_ value than the contralateral control region^14, 29^. With [^11^C]3Me4AP, this lesion was not visible on the PET images. The small size of this structure combined with the short half-life of carbon-11, as well as the moderate brain penetration and slow kinetics of [^11^C]3Me4AP resulted in noisy TAC, preventing us from being able to conclude that tracer uptake was enhanced in the craniotomy site (**Sup. Fig. 3**). While this finding does not completely exclude the possibility that [^11^C]3Me4AP may be able to detect large demyelinating lesions in the brain, it shows that the sensitivity of this tracer to this particular lesion was poor compared to [^18^F]3F4AP and [^11^C]3MeO4AP.

## Conclusion

In this study, we have developed a rapid and efficient radiochemical synthesis method of [^11^C]3Me4AP, a novel radiotracer for voltage-gated K^+^ channels based on the drug 4-aminopyridine. The synthesis proceeds *via* one-pot Pd(0)-Cu(I) mediated [^11^C]methylation of a stannyl precursor containing a free amino group. The labeling occurs under mild conditions and produces [^11^C]3Me4AP in good radiochemical yield, high molar activity, and excellent radiochemical purity (**Fig. 1**). The evaluation of [^11^C]3Me4AP in rodents showed moderate brain penetration, slow kinetics and an increase in brain signal upon coinjection of 3Me4AP, which is consistent with the mechanism that pharmacological doses of K^+^ channel blockers cause more channels to transition to the open/bindable state (**Fig. 2**). Further studies performed in NHPs confirmed the moderate brain penetration and slow kinetics (**Fig. 4**). Additionally, analyses of arterial blood collected in NHPs showed that [^11^C]3Me4AP is metabolically stable and mostly free in plasma (**Fig. 3**). The kinetics of [^11^C]3Me4AP in the different brain regions could be accurately described by a 1-tissue compartment model, which was used to calculate regional *V*_T_s (**Table 2**). Regional *V*_T_ estimates could also be accurately estimated using the MA1 graphical analysis method. Comparison of the regional *V*_T_ estimates with the V_T_s of its predecessors [^18^F]3F4AP and [^11^C]3MeO4AP suggested a binding affinity similar to [^11^C]3MeO4AP and higher than [^18^F]3F4AP (**Fig. 5**). Despite the good model fits, good overall test-retest reproducibility (<10% in average for both monkeys), high binding affinity and other favorable properties, the limited brain penetration and slower kinetics of [^11^C]3Me4AP resulted in a tracer with inferior performance than [^11^C]3MeO4AP and [^18^F]3F4AP. While this compound does not appear as promising as [^11^C]3MeO4AP and [^18^F]3F4AP for brain PET imaging, it may be useful as a therapeutic. Finally, given the high similarity in physicochemical properties (logP and p*K*_a_) and pharmacological properties (metabolic stability and binding affinity) of the therapeutic drug 4AP and 3Me4AP, the pharmacokinetic properties described here for 3Me4AP may provide a good approximation of the brain penetration and clearance of 4AP.

## Experimental Section

### Compliance

All rodent procedures were approved by the Animal Care and Use Committee at the Massachusetts General Hospital. All experiments involving non-human primates were performed in accordance with the U.S. Department of Agriculture (USDA) Animal Welfare Act and Animal Welfare Regulations (Animal Care Blue Book), Code of Federal Regulations (CFR), Title 9, Chapter 1, Subchapter A, Part 2, Subpart C, §2.31. 2017. All animal studies were conducted in compliance with the ARRIVE guidelines (Animal Research: Reporting in Vivo Experiments) for reporting animal experiments.

### Chemistry

#### General

Unless otherwise stated, all chemical reactions were carried out under an inert nitrogen atmosphere with dry solvents under anhydrous conditions. All chemicals were ordered from commercial suppliers and used without further purification. Reactions were monitored by thin-layer chromatography (TLC) carried out on 0.25 mm silica gel plates (60F-254, Merck) using UV light as the visualizing agent. Flash column chromatography employed SiliaFlash® P60 (40-60 μm, 230-400 mesh) silica gel (SiliCycle, Inc.) The ^1^H and ^13^C NMR spectra were collected on a 300 MHz Bruker spectrometer at 300 MHz and 75 MHz, respectively. Copies of the spectra can be found in the supporting information. All ^1^H-NMR data are reported in δ units, parts per million (ppm), and were calibrated relative to the signals for residual chloroform (7.26 ppm) in deuterochloroform (CDCl_3_). All ^13^C-NMR data are reported in ppm relative to CDCl_3_ (77.16 ppm) and were obtained with ^1^H decoupling unless otherwise stated. The following abbreviations or combinations thereof were used to explain the multiplicities: s = singlet, d = doublet, t = triplet, q = quartet, br = broad, m = multiplet. High-resolution mass spectra (HRMS) were recorded on a Thermo Scientific Dionex Ultimate 3000 UHPLC coupled to a Thermo Q Exactive Plus mass spectrometer system using ESI as the ionization approach.

The [^11^C]MeI was produced in the GE TRACERlab FX MeI synthesizer for the reported radiochemistry. Semipreparative HPLC separations were performed on Sykam S1122 Solvent Delivery System HPLC pump with the UV detector at 220 nm (for purification of [^11^C]3Me4AP). The purity of final compounds was verified by HPLC on a Thermo Scientific Dionex Ultimate 3000 UHPLC equipped with a waters XBridge BEH HILIC column (5 μm, 4.6 × 150 mm) and a diode array detector set at 254 nm.

#### Synthesis of precursor 3-(tributylstannyl)pyridin-4-amine (2)

To a solution of 3-iodo-4-aminopyridine (220 mg, 1.0 equiv, 1.0 mmol) in anhydrous THF (1.0 mL) at -40 °C, *t*-Butylmagnesium chloride solution (1.0 M in THF, 2.4 mL, 2.4 equiv, 2.4 mmol) was added dropwise while vigorously stirring. The white precipitate was formed immediately. After stirring the suspension at -40 °C for 30 min, the reaction was moved to a -78 °C cold bath. *t*-Butyllithium solution (1.7 M in pentane, 1.4 mL, 2.4 equiv, 2.4 mmol) was added dropwise, and the reaction mixture was allowed to warm up to room temperature by removing from the cold bath and stirring for 30 min. Then a solution of tributyltin chloride (0.4 mL, 1.5 equiv, 1.5 mmol) in THF (0.6 mL) was added to the reaction mixture at -78 °C, and the reaction mixture was warmed up slowly and stirred overnight. The reaction was quenched by sat. NH_4_Cl aqueous solution (6 mL) and extracted with AcOEt (10 mL × 3). The combined organic layers were dried over Na_2_SO_4_, and the solvent was removed *via* rotary evaporation. The residue was purified by flash chromatography with hexanes/AcOEt (2:1) as eluent giving the desired xx in 49% yield as light yellowish oil. The product was aliquoted for each vial containing 4 mg for further radiochemistry. R_f_ = 0.29 (hexanes/AcOEt = 2:1). ^**1**^**H NMR** (300 MHz, CDCl_3_): δ 8.21–8.14 (m, 2H), 6.50–6.43 (m, 1H), 4.07 (br, 2H), 1.67–1.44 (m, 6H), 1.39–1.25 (m, 6H), 1.23–1.00 (m, 6H), 0.88 (t, 9H); ^**13**^**C NMR** (75 MHz, CDCl_3_): δ 158.4 (*J*_*C-Sn*_ = 5.9 Hz), 156.8 (*J*_*C-Sn*_ = 15.5 Hz), 150.3, 119.1, 109.4, 29.2 (*J*_*C-Sn*_ = 29.5 Hz), 27.4 (*J*_*C-Sn*_ = 10.2 Hz), 13.7, 9.8 (*J*_*C-Sn*_ = 168.2 Hz); **ESI-HRMS** (m/z): [M+H]^+^ calc’d for C_17_H_33_N_2_Sn^+^: 385.1660; found: 385.1659.

#### Radiosynthesis of [^11^C]3Me4AP (3)

In an oven-dried Thermo 3 to 5 mL V-vial, equipped with a magnetic stir bar, was weighted tris(dibenzylideneacetone)dipalladium(0) (3.5 mg, 0.4 equiv, 3.8 μmol), tri(*o*-tolyl)phosphine (3.5 mg, 1.1 equiv, 11.5 μmol), cesium fluoride (2.2 mg), and copper(II) bromide (2.8 mg). The aliquoted precursor xx (4 mg, 1.0 equiv, 10.4 μmol) was dissolved with anhydrous N-methyl-2-pyrrolidone (NMP, 0.2 mL) and transferred to the V-vial reactor. The complete transfer of the precursor is by rinsing with an additional 0.15 mL of NMP. The reaction vial was sealed with the cap under the inert atmosphere and pre-stirred for 5 minutes. Then [^11^C]MeI was bubbled into the reaction solution and allowed to react at room temperature for 15 minutes. The reaction mixture was diluted with 3.5 mL of mobile phase (described below) and purified using HPLC (SiliChrom HILIC column, 5μm, 10 × 150 mm; 25% Acetone in 75% ammonium bicarbonate aqueous (10 mM, pH = 8); flow 5 mL/min; t_**R**_ ∼ 10.5 min, collecting from ∼9.7 min to ∼11.4 min). The non-decay-corrected RCC was determined to be 89±5% from the reaction crude by HPLC (70% 10mM NH_4_HCO_3_ aqueous and 30% acetonitrile). Compound identity was confirmed by coinjection with nonradioactive reference standard on HPLC (30% 10mM NH_4_HCO_3_ aqueous and 70% acetonitrile, t_**R**_ ∼ 7.2 min on radioactive spectra). See molar activity calculation curve and NMP amount determination in the SI. The collected solution was diluted with saline to achieve a solution containing no more than 10% acetone by volume for the animal studies.

### Rats Imaging Studies and Measurement of Brain SUV ex vivo

#### Animals

6–8-week-old male and female Sprague Dawley rats were used.

#### Imaging Studies and Measurement of Brain SUV ex

Both naïve female and male rats were imaged on a Sedecal SuperArgus PET/CT scanner. Rats were injected via the tail vein with the radioligand only (approximately 0.4 mCi in 1 mL solution) or radioligand (same) plus 5 mg/kg of cold 3Me4AP and scanned for 60 mins under anesthesia (2% isoflurane with an oxygen flow of 2.0 L/min). After completing the scan, the animals were euthanized, their blood was collected by cardiac puncture and their brains were harvested. Blood and brain samples were weighed, and the radioactive concentrations were measured using a single well gamma counter. Four rats (2 male, 2 female) were used in imaging experiments. Fifteen rats (11 male, 4 female) were used in gamma counter experiments.

### NHP PET imaging studies

#### Animals

Two male adult rhesus macaques (monkey 1 and monkey 2) were used in the study (ages 10 and 14 years old, respectively). Their body weights on the day of imaging were 14.2 kg for monkey 1 and 17.5 kg for monkey 2. As previously reported, one of the animals (monkey 2) had sustained an accidental injury during a craniotomy procedure 4 years prior to imaging^14, 16^.

#### Animal Preparation

Prior to each imaging session, animals were sedated with ketamine/xylazine (10/0.5 mg/kg IM) and were intubated for maintenance anesthesia with isoflurane (1-2% in 100% O_2_). A venous catheter was placed in the saphenous vein for radiotracer injection, and an arterial catheter was placed in the posterior tibial artery for blood sampling. The animal was positioned on a heating pad on the bed of the scanner for the duration of the study.

#### Imaging procedure

A three-dimensional T1-weighted magnetization-prepared rapid gradient-echo (MEMPRAGE) was acquired in each animal to provide an anatomical reference with acquisition parameters as described previously^14, 16^.

The two rhesus macaques each underwent two baseline dynamic PET scans on a Discovery MI (GE Healthcare) PET/CT scanner. The two scans were acquired on the same day for Monkey 1 and were separated by 1 month for monkey 2. A CT scan was performed before each PET acquisition for the purpose of correcting the PET images for photon attenuation. Emission PET data were acquired for 180 min in 3D list mode and data collection was started right before [^11^C]3Me4AP injection. [^11^C]3Me4AP was administered as a 3-min infusion and was followed by a 3-min infusion of a saline flush. Radiotracer and saline flush were injected using two Medfusion 3500 syringe pumps. At the time of injection, [^11^C]3Me4AP doses were 304.6 and 314.2 MBq for Monkey 1, and 330.1 and 291.5 MBq for Monkey 2. Corresponding molar activities (Am) of [^11^C]3Me4AP were 90.3 and 247.5 GBq/μmol for Monkey 1, and 162.8 and 176.1 GBq/μmol for Monkey 2 at the time of injection. PET images were reconstructed with a 3D time-of-flight iterative reconstruction algorithm using 3 iterations and 34 subsets with corrections for scatter, attenuation, deadtime, random coincident events, and scanner normalization. Dynamic images were reconstructed with the following time frames: 6 × 10, 8 × 15, 6 × 30, 8 × 60, 8 × 120 s and remaining were 300 s frames. Final PET images had voxel dimensions of 256×256×89 with voxel sizes of 1.17×1.17×2.8mm^3^.

#### Arterial Blood and Radiometabolite Analysis

Blood samples were manually collected throughout the PET acquisition in order to measure the radioactivity time courses in whole-blood (WB) and plasma (PL) and to characterize [^11^C]3Me4AP in-vivo stability by radio HPLC. The procedures adopted to measure WB and PL curves in kBq/mL and for subsequent transformation into SUV units were similar to those previously described in Guehl *et al*. ^16^. [^11^C]3Me4AP radiochromatograms and corresponding parent fractions were measured for plasma samples drawn at 3, 5, 10, 15, 30, 60, 90 and 120 min after radiotracer injection. The plasma samples were further filtrated with 10K filter (Amicon™ Ultracel-10 regenerated cellulose membrane, 0.5 mL sample volume, Centrifugal Filter Unit) by centrifugation at 21000 RCF (8 °C) for 15min. The filtrates were injected into HPLC (Agilent 1260 Infinity II, along with Eckert & Ziegler FlowCount radioisotope detector) through a C18 column (XBridge, BEH, 130 Å, 3.5 μm, 4.6 × 100 mm) with a vanguard cartridge (XBridge, BEH, 130 Å, 3.5 μm, 3.9 × 5 mm) using 12 mM Na_2_HPO_4_ /8 mM NaOH aqueous solution (pH=12) : MeCN = 96:4 as mobile phase. For each study, the plasma free fraction (*f*_*p*_) was measured in triplicate as previously described ^14^.

#### PET Data processing and image quantification

Dynamic PET images were registered to the corresponding MEMPRAGE images via a rigid body transformation and the structural MEMPRAGE images were aligned to the National Institute of Mental Health macaque template (NMT) ^**30**^ via affine followed by non-linear transformations. The inverse transformation matrices were calculated and applied to the NMT atlases for extraction of brain time activity curves (TACs) in the native PET space. Registration procedures were performed using FLIRT and FNIRT from the FMRIB Software Library ^**31**^. TACs were extracted for 10 brain regions (occipital cortex, parietal cortex, temporal cortex, frontal cortex, hippocampus, amygdala, striatum, thalamus, white matter, and whole cerebellum) in Bq/cc and were subsequently transformed in SUV unit.

TACs were analyzed by compartmental modeling using one-tissue (1T) and two-tissue (2T) models and the metabolite-corrected input function. The total volume of distribution (V_T_) was the primary outcome measure used in this work to quantify [^11^C]3Me4AP uptake and was calculated from the model microparameters. In addition, the Logan graphical method ^**32**^ and the multilinear analysis 1 (MA1) method^**33**^ were tested as simplified methods for the direct estimation of *V*_*T*_. The consensus nomenclature for in vivo imaging of reversibly binding radioligands described in Innis et al. ^**34**^ was followed in this work.

#### Quantitative comparison of in vivo binding properties between [^11^C]3Me4AP, [^18^F]3F4AP and [^11^C]3MeO4AP

In each animal, we performed a direct comparison of the *in vivo* affinity and specific binding ratios of [^11^C]3Me4AP *vs*. [^18^F]3F4AP and [^11^C]3Me4AP *vs*. [^11^C]3MeO4AP using regional *V*_*T*_ values *via* linear regression following the method reported by Guo et al. ^16, 28^ that we previously used when we reported the comparison between [^11^C]3MeO4AP and [^18^F]3F4AP ^16^. Briefly, in this graphical method, *V*_*T*_ of [^11^C]3Me4AP and *V*_*T*_ of [^18^F]3F4AP are directly related to the *in vivo* affinity and specific binding ratios as shown in eq 1 below,

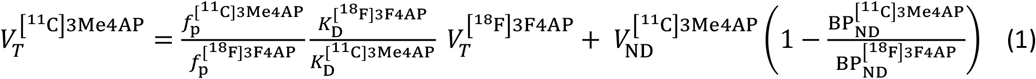

Where 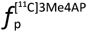 and 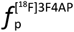 are the plasma free fractions, 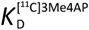 and 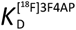 are the equilibrium dissociation constants, 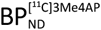 and 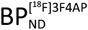 are the binding potentials (the ratio at the equilibrium of specifically bound radioligand to that of nondisplaceable radioligand in tissue) for [^11^C]3Me4AP and [^18^F]3F4AP, respectively, and 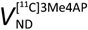 is the nondisplaceable volumes of distribution of [^11^C]3Me4AP assumed to be homogeneous across brain regions. By plotting of *V*_*T*_ of one tracer against *V*_*T*_ of the other tracer across several brain regions with different levels of binding, this graphical method allows to (1) determine whether the two tracers bind to the same target (assessed by the linearity or lack of linearity of the regression) and (2) identify which compound presents the higher signal-to-noise ratio (assessed from the sign of the y-intercept). In addition, since measurements of 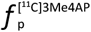 and 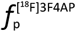 are available and both [^11^C]3Me4AP and [^18^F]3F4AP enter the brain by passive diffusion, the slope of the linear regression provides information on the *in vivo* affinity ratio ^28^. In each animal, the comparison was performed on the average *V*_*T*_ values of the radiotracers calculated from the two baseline scans. Regional [^18^F]3F4AP *V*_*T*_ data were obtained from Guehl et al. ^14^ (where monkeys 3 and 4 correspond, respectively, to monkeys 1 and 2 in the present work). Paired comparisons were plotted for the regional *V*_*T*_ estimates in the occipital cortex, parietal cortex, temporal cortex, frontal cortex, hippocampus, amygdala, striatum, thalamus, white matter, and whole cerebellum. A similar comparison between [^11^C]3Me4AP and [^11^C]3MeO4AP was performed.

#### Statistical Analysis

All data are expressed as the mean value ± one standard deviation (SD). The Akaike information criterion (AIC) was used to assess the relative goodness of fit between compartment models^26^. The most statistically appropriate model is the one producing the smallest AIC values. The linear correlation between *V*_*T*_ of [^11^C]3Me4AP and *V*_*T*_ of [^18^F]3F4AP (eq 1) was assessed using Pearson’s correlation coefficient r, and the t distribution of the Fisher transformation was used to generate p values for linear regressions and intercept. A p-value of 0.05 or less was considered statistically significant. Test−retest variability (TRT in %) of *V*_*T*_ estimates was calculated from the two baseline scans acquired on two days 1 month apart in between in monkey 2 as TRT(%) = 100 × 2 x abs(*V*_*T*,baseline1_ − *V*_*T,baseline2*_)/(*V*_*T,baseline1*_+ *V*_*T,baseline2*_). The same equation was also used to calculate the reproducibility of *V*_*T*_ measurements in monkey 1 from the baseline scans that were acquired one month apart on the same day.

## Acknowledgments

The authors thank David F. Lee, Jr., Dr. John A. Correia and Dr. Hamid Sabet for providing the carbon-11 for the radiotracer synthesis. We thank Jennifer X. Wang for the high-resolution mass spectra measurement and data analysis.

## Author contributions

YS developed the radiosynthesis method, produced the tracer and performed quality control. YS, YPZ, VB and KT performed PET imaging in rats, rat tissue dissections, gamma counting, and data analysis of the rodent data. NJG, GEF and MDN performed PET imaging in NHPs and analyzed the data. YS, MD and SHM processed and analyzed the NHP blood samples. YS, NJG and PB wrote the manuscript, which was reviewed and approved by all the authors.

## Funding

This study was supported by NIH R01NS114066 (PB), P41EB022544 (GEF), S10OD026987 (MDN, GEF).

## Competing interest

PB has a financial interest in Fuzionaire Diagnostics and the University of Chicago. PB is the inventor of a PET imaging agent owned by the University of Chicago and licensed to Fuzionaire Diagnostics. Dr. Brugarolas’s interests were reviewed and are managed by MGH and Mass General Brigham in accordance with their conflict-of-interest policies. The other authors declare no conflict of interests.

