## Supplemental information for "Radiochemical Synthesis and Evaluation of 3-[^11^C]Methyl-4-aminopyridine in Rodents and Non-Human Primates for Imaging Potassium Channels in the CNS"

**{$^{1}$H NMR} (up) and {$^{13}$C NMR} (bottom) spectra of 3-(tributylstannyl)pyridin-4-amine:**

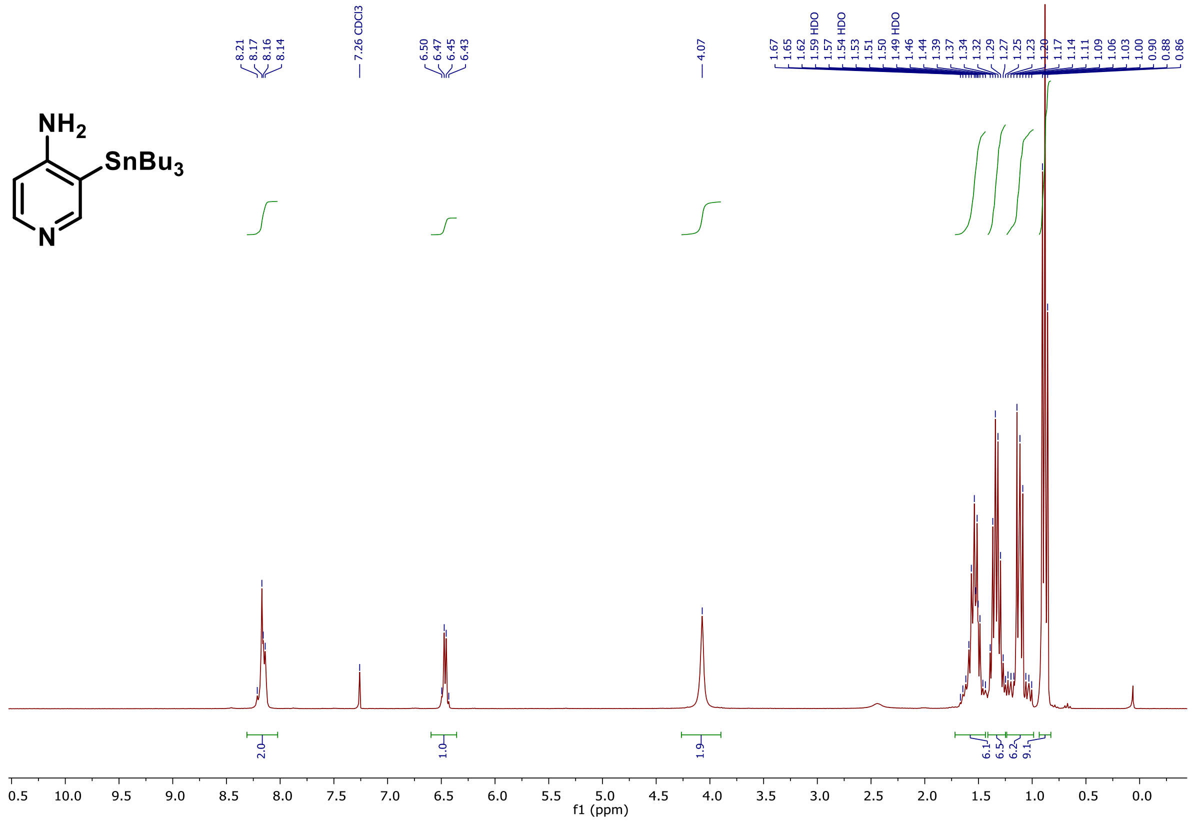

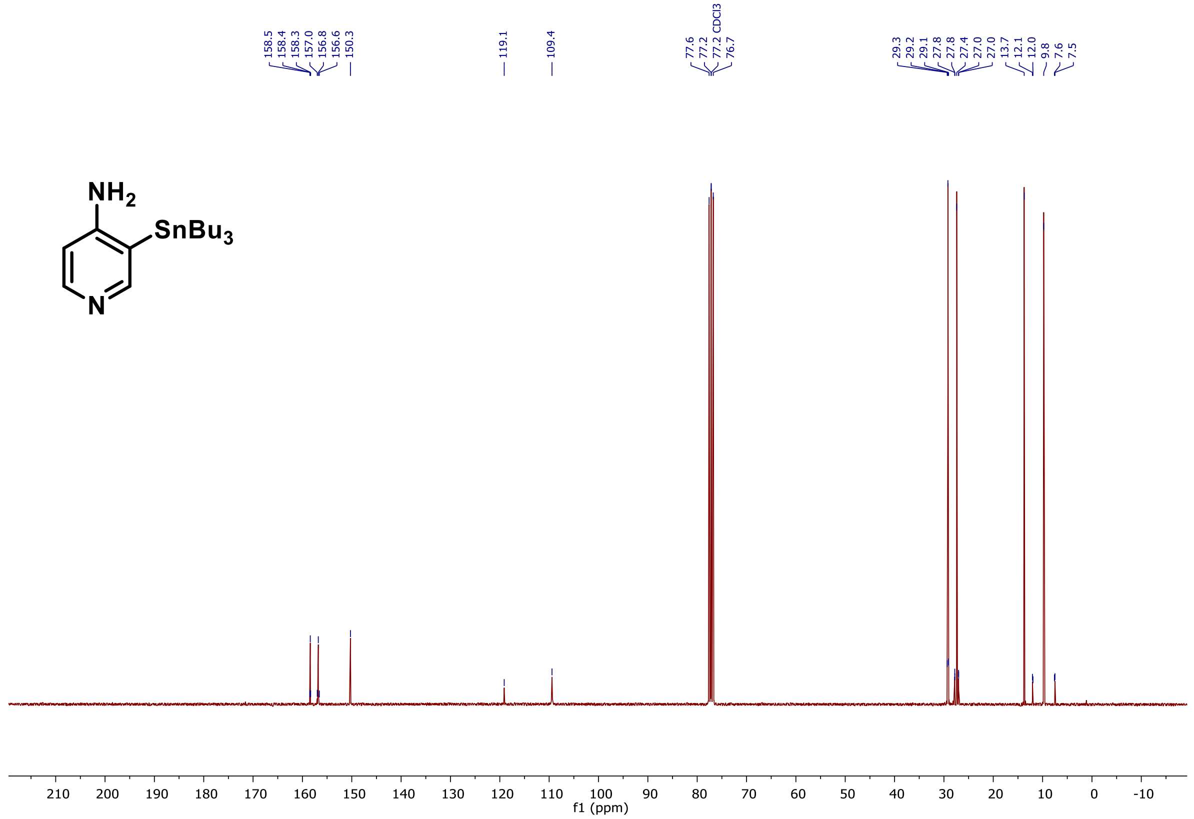

**The molar activity calculation curve** was determined by injecting serial concentrations of 3Me4AP solution in the same volume into HPLC to get the corresponding area under the curve at 254 nm.

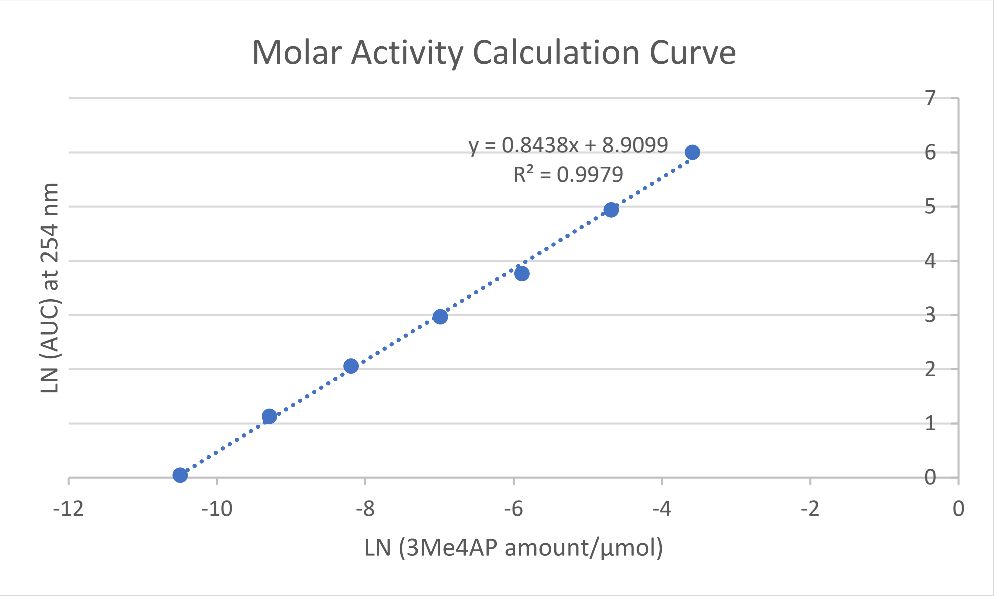

**NMP amount in the [$^{11}$C]3Me4AP solution determination**: NMP calculation curve was determined by injecting serial amounts of NMP into HPLC to get the corresponding area under the curve at 210 nm with waters XBridge C18 column (3.55 µm, 4.6 × 150 mm) and gradient 3 to 30% acetonitrile in 10mM $NH_4HCO_3$ aqueous solution as mobile phase.

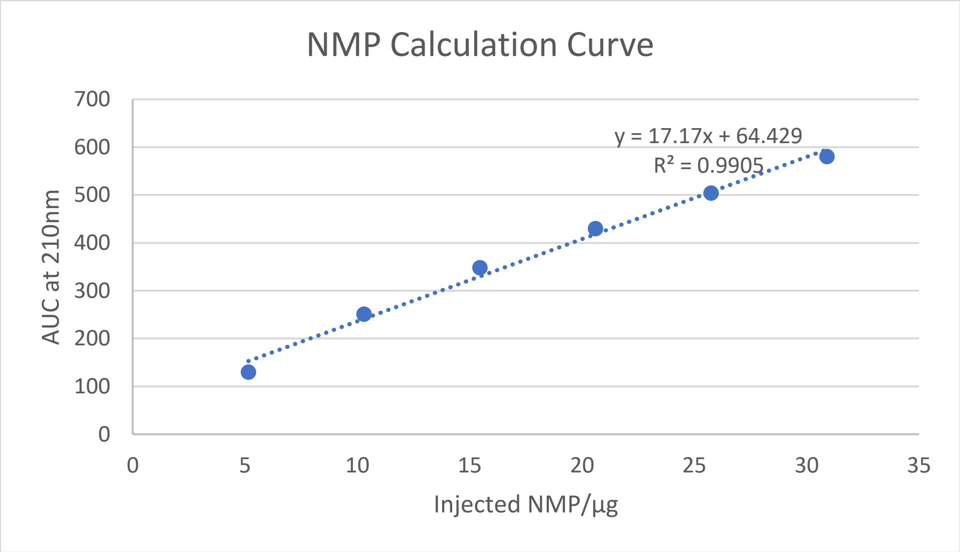

Then the final [11C]3Me4AP solution in the semiprep HPLC mobile phase was injected into the HPLC and calculated finding that the concentration of NMP is 43±8 ppm, which means the absolute NMP amount in 10 mL solution is 0.43±0.08 mg.

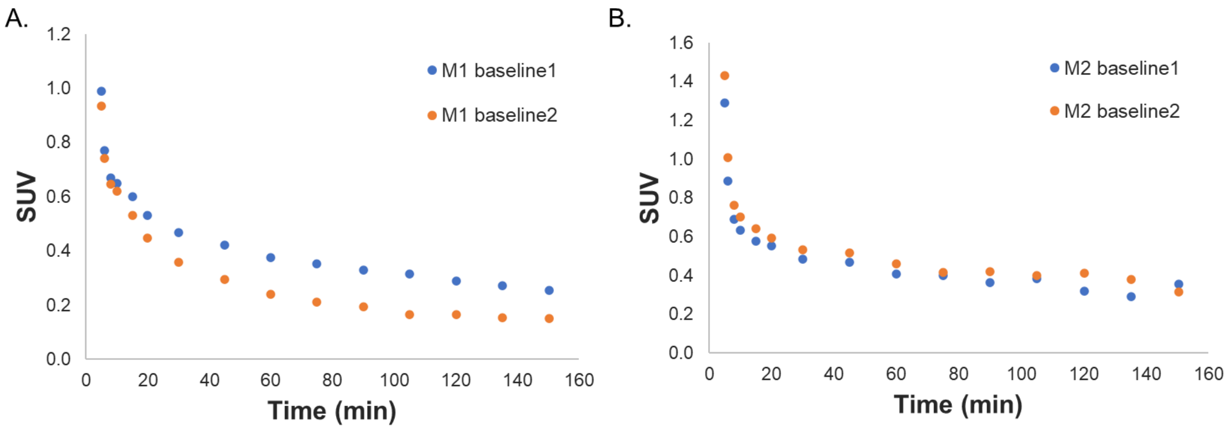

**Figure 1.** [$^{11}$C]3Me4AP concentration in plasma of monkey 1 and monkey 2: time–activity curves from 5 min to 150 min.

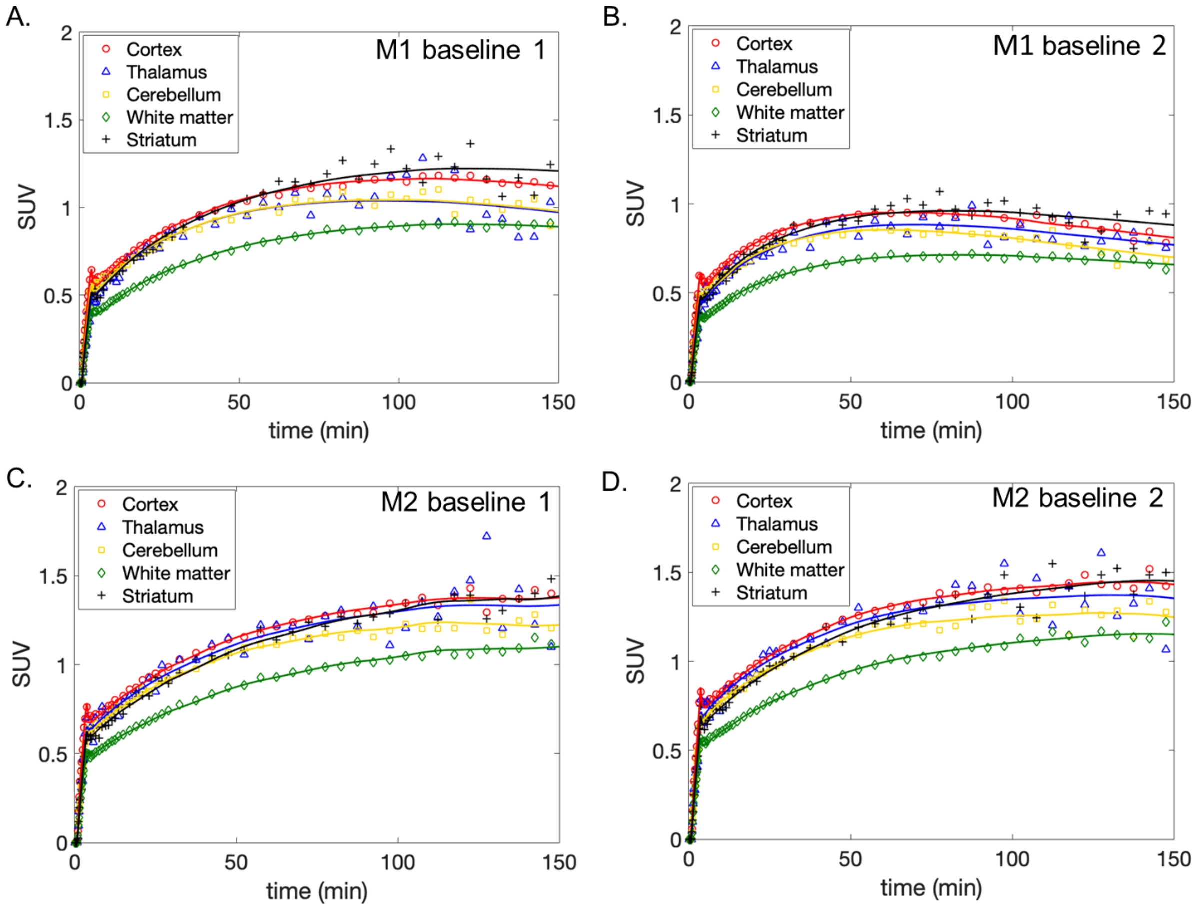

**Figure 2.** NHP time–activity curves of selected brain regions in monkey 1 and monkey 2 with two replicated baseline scans.

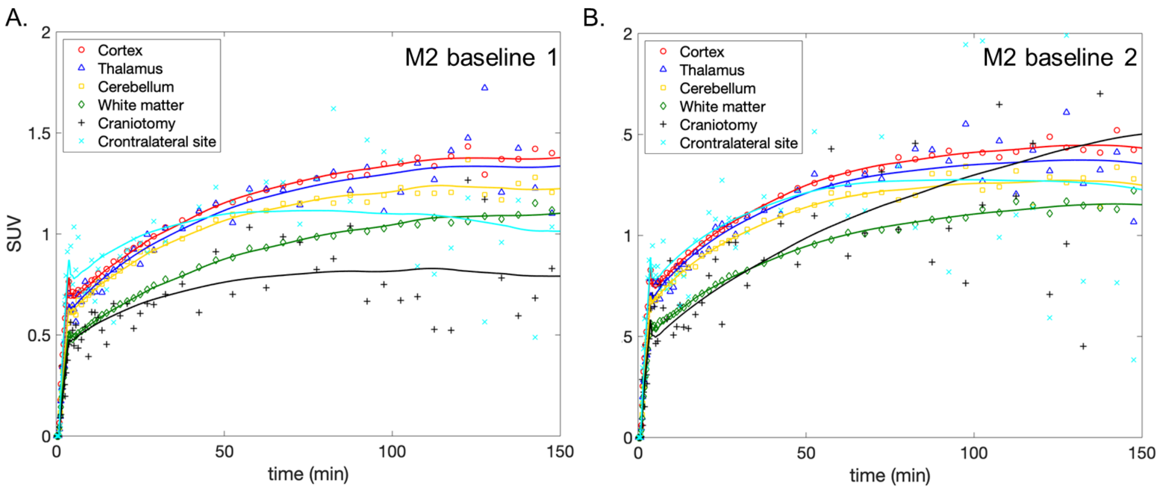

**Figure 3.** NHP time–activity curves including the lesion in light blue of monkey 2 with two replicated baseline scans.
